# Cramming Protein Language Model Training in 24 GPU Hours

**DOI:** 10.1101/2024.05.14.594108

**Authors:** Nathan C. Frey, Taylor Joren, Aya Abdelsalam Ismail, Allen Goodman, Richard Bonneau, Kyunghyun Cho, Vladimir Gligorijević

**Affiliations:** Prescient Design, Genentech

## Abstract

Protein language models (pLMs) are ubiquitous across biological machine learning research, but state-of-the-art models like ESM2 take hundreds of thousands of GPU hours to pre-train on the vast protein universe. Resource requirements for scaling up pLMs prevent fundamental investigations into how optimal modeling choices might differ from those used in natural language. Here, we define a “cramming” challenge for pLMs and train performant models in 24 hours on a single GPU. By re-examining many aspects of pLM training, we are able to train a 67 million parameter model in a single day that achieves comparable performance on downstream protein fitness landscape inference tasks to ESM-3B, a model trained for over 15, 000*×* more GPU hours than ours. We open source our library^1^ for training and inference, **LBSTER**: **L**anguage models for **B**iological **S**equence **T**ransformation and **E**volutionary **R**epresentation.

## 1 Introduction

Protein Language Models (pLMs) are a powerful framework for representation learning across the large, diverse protein universe that have become critical components for predicting protein structure and function (Lin et al., 2023; Chen et al., 2023; Elnaggar et al., 2022; Xu et al., 2023). Current SOTA pLMs require enormous compute budgets to scale up model size and training time. Expensive and time-consuming pre-training, however, makes it infeasible for most practitioners to rapidly experiment and understand pLM performance. To enable greater exploration, rapid pre-training of performant pLMs is essential. To this end, in this paper we introduce a “cramming” challenge for pLMs - where the objective is to train a pLM in a single day on a single GPU - and we propose new architectural and training choices that maximize performance of “scaled down” pLMs. These “crammed” pLMs perform competitively with SOTA ESM2 models on downstream functional prediction tasks from FLIP (Dallago et al., 2021) and protein-protein interaction (PPI) (Mei & Zhang, 2019) classification. We envision that others will build on our work to propose even better cramming strategies for pLMs, and that our work will enable more rapid progress on pre-trained models for biology. All training and model code and open-source model weights and dataset splits are available in our library, **LOBSTER**: **L**anguage m**O**dels for **B**iological **S**equence **T**ransformation and **E**volutionary **R**epresentation.

## 2 Training a protein language model on a single GPU in a single day

### 2.1 Defining the cramming challenge setting

We first define the settings for our challenge, which are mostly borrowed from (Geiping & Goldstein, 2023). The rules for pLM cramming are:

- A transformer-based language model is trained from scratch with a masked-language modeling objective.
- Training may not exceed 24 hours on a single GPU.
- No existing pre-trained models are used at any point.
- The training, validation, and test data splits are from UniRef50 and these are pre-specified. The training data can be sampled in any way that does not involve a pre-trained model, hence speedups may be achieved by careful choices of how and when to sample training data.
- The downloading of raw data in FASTA format is exempt from the overall compute budget. All preparation of raw FASTA inputs for training (e.g., tokenization, filtering, sorting, etc.) happens on-the-fly during training and is included in the training budget.
- Downstream performance is evaluated on tasks from the FLIP (Dallago et al., 2021) and PPI (Mei & Zhang, 2019) benchmarks. Hyperparameters are set globally for all downstream tasks. Any aggregation method can be used to pool embeddings from the crammed model and any architecture can be used for the prediction head for downstream tasks, but these choices must be set globally for all downstream tasks. Downstream finetuning is not included in the 24 GPU hour cramming compute budget.

Our goals and therefore experimental settings are different from those in (Geiping & Goldstein, 2023). The goals of pLM cramming are to 1) enable rapid experimentation and training of “production” pLMs to re-examine fundamental assumptions about how language modeling is applied to biological sequence data; 2) apply interpretability techniques that require intervening on model training (and therefore the ability to retrain models often); and 3) better understand “scaling down” and what really matters for downstream pLM performance (architecture, model size, optimizer, dataset construction, etc.). With these goals in mind, we have constructed the pLM cramming rules to make our setup as simple as possible to replicate (fixing the dataset and train/val/test splits, no data pre-processing steps outside of training, global hyperparameters for downstream evaluation, etc.).

Historically, pLM architectures and training setups hew very close to the original work of Devlin et al. (2018), going so far as to keep everything identical except the vocabulary and dataset. Following Geiping & Goldstein (2023), we seek to maximize per-token efficiency of training, and propose architectural and training modifications intended to achieve this goal. The scaling literature (Geiping & Goldstein, 2023; Kaplan et al., 2020) indicates that per-token efficiency depends strongly on model size but is largely invariant to model shape. Smaller models learn less efficiently, so the most impactful changes speed up gradient computation for a fixed model size.

All hyperparameters, pooling, and architectures choices must be set globally for all downstream tasks. To ensure that downstream finetuning remains a negligible compute cost compared to the cramming pre-training, we limit finetuning on a single task to 10% (2.4 hours) of the overall cramming compute budget. We also evaluate finetuning performance with no time limit to measure the performance of large models, which train more slowly but may also reach better performance if given unlimited compute. These limitations are in line with our goals of enabling rapid experimentation, while also providing flexibility to find the best cramming + finetuning setup for downstream performance.

All the experiments reported in this paper are conducted on NVIDIA A100-SXM4-80GB GPUs, with Python 3.10.9, pytorch 2.0.1, cudatoolkit 11.7, transformers 4.30.2, and lightning 1.9.5.

### 2.2 pLM modifications

#### Architecture modifications

We adapt the HuggingFace implementation of the ESM2 architecture (Lin et al., 2023) as a starting point for cramming. To maximize per-token training efficiency, we remove all query, key, and value biases in all attention blocks (Dayma et al., 2021). This reduces computation without greatly affecting overall parameter account. Similarly, we remove all bias terms in intermediate linear layers (Dayma et al., 2021).

#### Training modifications

To achieve a large effective batch size despite the cramming constraints, we accumulate gradients and perform updates every 16 forward/backward passes. We use a batch size of 128 and a maximum length of 512 (large enough to accommodate most single proteins in the training dataset), for a total effective batch size of 2048 sequences or 1,048,576 tokens. We adjust the masking rate from the standard 15% used in the BERT (Devlin et al., 2018) and ESM2 (Lin et al., 2023) setups to 25%, as 15% leads to awkward tensor shapes and 25% of the sequence length is 128 = 2^7^. We also hypothesize that, due to evolutionary relationships among protein sequences, 15% is far too low a masking rate and we can train better pLMs more efficiently by making the pre-training denoising task more difficult. We use the AdamW (Loshchilov & Hutter, 2017) optimizer with *β*_1_ = 0.99, *β*_2_ = 0.98, and *ε* = 10^*−*12^. A gradient clipping value of 0.5 is used to stabilize training. Training is performed with automated mixed precision (Micikevicius et al., 2017).

We find that the learning rate and learning rate schedule are by far the most important hyperparameters for pLM cramming. We thoroughly ablate these hyperparameters and present the results below in Section 4. Tuning the learning rate schedule to achieve the maximum learning rate possible without causing training instabilities is vital to pLM cramming performance. To anneal the learning rate to near zero within the allotted 24 GPU hours, we first estimate the total training budget and set the maximum number of steps to 50,000. In our experiments, we found the best performance with a maximum learning rate of 1 *×* 10^*−*3^ and a linear learning rate decay with a warmup period of 1,000 steps. This corresponds to a fast warmup and slow cooldown, which interestingly is the exact opposite of the optimal learning rate schedule found in Geiping & Goldstein (2023).

#### Opportunities for further optimization

There are a few obvious opportunities for further training efficiency increases that we leave for future work. We perform validation loss checks throughout training to monitor training performance and stability; these could be disabled to avoid the unnecessary compute cost. There are other logging and profiling capabilities in Lightning that can be disabled as well. Using 8-bit floating point mixed precision training and other recent advances in efficient transformer training are also promising avenues for future work.

## 3 Related work

### 3.1 Efficient transformers

The most closely related work to ours is Geiping & Goldstein (2023). However, due to the fundamental differences between biological sequence data and natural language noted above, the goals, implementations, and results of our work differ substantially. Izsak et al. (2021) trained BERT models on a full server node of 8 GPUs in a single day. Much recent work is focused on improving the efficiency of training transformers (Treviso et al., 2023), but most architectural changes do not show persistent performance improvements over many orders of magnitude of model and dataset sizes (Kaplan et al., 2020).

### 3.2 Efficient protein language models

Elnaggar et al. (2023) sought to achieve state-of-the-art pLM performance while reducing the overall model size compared to ESM2 (Lin et al., 2023), by ablating architectural and dataset construction choices, but with no restrictions on compute budget. Serrano et al. (2023) introduced “Small-Scale Protein Language Model (SS-pLM)”, a 14.8M parameter model for rapid experimentation. We do not put any restrictions on model size, and instead focus on architectural and training choices that maximize token throughput and speed up convergence. Yang et al. (2022) did away with transformers altogether and showed that pre-trained convolution-based architectures are significantly cheaper and competitive with transformer-based pLMs. Since the first appearance of our work, Li et al. (2024) showed in a thorough evaluation of 370 pLM transfer learning experiments that almost all downstream task evaluations benefit from using pLM representations, but performance on the majority of public benchmark tasks does not scale with pre-training. This reliance on low-level features learned early in pre-training agrees with our findings.

## 4 Experiments

### 4.1 Learning rate dynamics

The results of the hyperparameter sweep over learning rates and number of warmup steps are presented in Table 1. We sweep over a range of learning rates *∈* [1 *×* 10^*−*2^, 4 *×* 10^*−*4^] and number of warmup steps *∈* [100, 40000]. We find that the choice of learning rate and warmup steps has a huge impact on the validation perplexity, which ranges from 13.72 for the best model and 20.49 for the worst with a vocabulary size of 33. The optimal hyperparameter choices allow for a maximally high learning rate, with a schedule that prevents training instabilities and anneals the learning rate close to zero by the end of training. In our experiments, the best model reaches a maximum learning rate of 0.001 after 1000 warmup steps, and then does a slow annealing of the learning rate over the remaining 49000 steps.

**Table 1:**
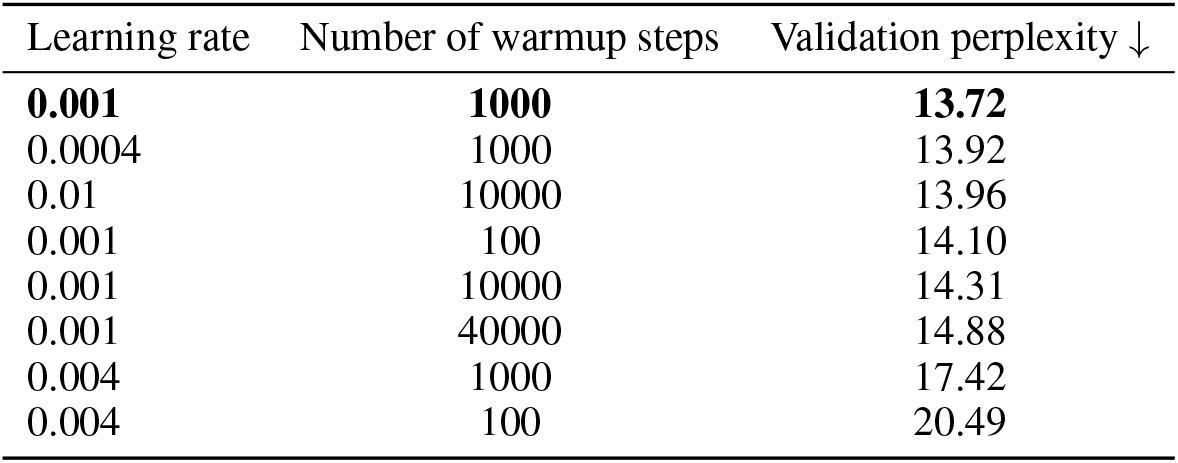
pLM cramming learning dynamics.

### 4.2 Downstream task evaluation

The results of downstream task evaluation are shown in Tables 2, 3 and 4. We evaluate our crammed models on four tasks including three protein fitness landscape inference tasks from the FLIP (Dallago et al., 2021) benchmark – GB1, AAV, and Meltome – and one protein-protein interaction (PPI) task from Mei & Zhang (2019). The FLIP benchmark contains many train/test splits based on edit distance and sequence similarity to provide a detailed evaluation of a model’s ability to “generalize” in different realistic protein engineering settings. First, as is typical in the machine learning literature, we evaluate downstream performance using IID splits for the GB1 and AAV tasks. As noted in (Dallago et al., 2021), random splits are not particularly interesting to biologists, but they greatly simplify evaluation. We take the train/test splits from Dallago et al. (2021), and as in that work, we randomly sample 10% of the training set as the validation set. We also evaluate OOD generalization using the 2-vs-rest splits for both GB1 and AAV. The Meltome dataset does not provide an IID split, so we use only the “mixed split” based on cluster components.

**Table 2:** Downstream task evaluation with a 10% cramming time limit. The FLIP (Dallago et al., 2021) tasks (GB1, AAV, and Meltome) results are reported in Spearman correlation and PPI are reported in AUPRC.

| Model | GB1 | AAV | Meltome | PPI |
| --- | --- | --- | --- | --- |
| **Crammed pLM-67M (Ours)** | **0.53** | **0.76** | **0.34** | **0.78** |
| ESM2-8M | 0.59 | 0.81 | 0.42 | 0.86 |
| ESM2-150M | 0.55 | 0.78 | 0.29 | 0.88 |
| ESM2-3B | 0.40 | 0.62 | 0.20 | 0.85 |

**Table 3:** Downstream task evaluation with no time limit. The FLIP (Dallago et al., 2021) tasks (GB1, AAV, and Meltome) results are reported in Spearman correlation and PPI are reported in AUPRC.

| Model | GB1 | AAV | Meltome | PPI |
| --- | --- | --- | --- | --- |
| Crammed pLM-67M (Ours) | 0.63 | 0.79 | 0.51 | 0.78 |
| ESM2-8M | 0.59 | 0.83 | 0.59 | 0.86 |
| ESM2-150M | 0.58 | 0.82 | 0.63 | 0.88 |
| ESM2-3B | 0.66 | 0.81 | 0.54 | 0.88 |

**Table 4:** FLIP downstream task evaluation (sp) with no time limit on OOD test splits. Each model is compared with its randomly-initialized baseline to highlight gains from pre-training.

| Model | Pre-trained |  |  | Baseline (no pre-training) |  |  |
| --- | --- | --- | --- | --- | --- | --- |
|  | GB1 | AAV | Meltome | GB1 | AAV | Meltome |
| Crammed pLM-67M (Ours) | 0.42 | 0.12 | 0.41 | 0.33 | -0.03 | 0.48 |
| ESM2-8M | 0.16 | 0.29 | 0.29 | -0.02 | -0.10 | -0.19 |
| ESM2-150M | 0.16 | 0.38 | 0.44 | 0.17 | -0.13 | -0.21 |
| ESM2-3B | 0.19 | 0.20 | 0.36 | 0.30 | -0.10 | -0.23 |

The PPI benchmark is to classify pairs of protein sequences as interacting or non-interacting. We use the Neglog dataset (Mei & Zhang, 2019), which consists of positive, interacting pairs as well as negative, non-interacting pairs augmented from Negatome 2.0 (Blohm et al.). We create an IID split by randomly sampling 10% as the test set with 70% used for training and 20% for validation.

In Tables 2 and 3, we report the validation set performance, as we have performed no hyperparameter tuning for downstream evaluation. We additionally report the test set performance in Table 4 for all OOD splits. We freeze the encoders and train a simple two layer multi-layer perceptron (MLP) with a feed-forward dimension of 256 for each task and a constant learning rate of 4 *×* 10^*−*5^ and batch size of 128. Token embeddings are aggregated using mean pooling prior to the MLP.

We consider three baselines, which are ESM2 (Lin et al., 2023) models of size 8M, 150M, and 3B parameters. These models are trained on over 60M unique protein sequences from UniRef50 and UniRef90, with an effective batch size of 2M tokens. The learning rate was warmed up over 2,000 steps to a peak value of 4 *×* 10^*−*4^ and then linearly decayed to 4 *×* 10^*−*5^ over 90% of the training duration, for a total of 500K training steps. Crucially, the 3B parameter model was trained on *512 NVIDIA V100 GPUs over 30 days*, or 368,640 GPU hours. In contrast, our crammed models were trained in 24 GPU hours, representing 0.0065% of the total training time of ESM2-3B, or a 15, 000*×* speedup.

In Table 2, we show results for all four tasks where finetuning is limited to 10% of the cramming time limit (2.4 GPU hours). In this regime, we find that downstream performance is inversely correlated with model size. Smaller models train faster and in the 2.4 GPU hour time limit, model capacity (size) does not compensate for this. In Table 3 we show results for finetuning with no time limit; models are trained to convergence with early stopping to prevent overfitting. Our crammed model achieves comparable performance on the FLIP and PPI downstream task evaluations to the significantly larger ESM2 models, but finetuning is completed in a small fraction of the time it takes for larger models.

Table 4 reports results on the OOD test splits. The crammed model outperforms ESM baselines on the GB1 and Meltome tasks, suggesting that 24 hrs of pre-training can effectively produce representations that generalize to OOD data. We additionally compare each pre-trained encoder to its randomly initialized baseline to highlight the gains only explained by pre-training. In several cases, the pre-trained model does worse than its randomly initialized counterpart, likely because the trainable MLP is driving performance more than the pre-training, a phenomenon seen in both crammed and non-crammed pLMs. Many models do not generalize OOD regardless of the amount of pre-training time. We speculate the global fine-tuning strategy chosen, which uses mean pooling, is suboptimal for representing proteins for OOD landscape prediction tasks. We leave it to future work to more thoroughly study global fine-tuning strategies within the rules of the cramming challenge.

## 5 Conclusions

In this paper we introduced the “cramming” challenge for protein language models - wherein the challenge is to train a performant pLM in 24 hours on a single GPU. To cram pLMs, we re-examine many parts of the original BERT model and training setup that, until now, pLMs have largely followed to the letter. By making architectural and training modifications to maximize per-token training efficiency, we are able to efficiently train pLMs in 24 GPU hours. The peak learning rate and learning rate schedule (number of warmup steps) are found to be by far the most important hyperparameters to minimize validation perplexity during pre-training. We evaluate our best crammed model against three ESM2 baselines on three protein fitness landscape inference tasks and a protein-protein interaction task and find that, using only 0.0065% of the total training time of ESM2-3B, our crammed model is largely competitive with the SOTA ESM2 baselines. We expect more difficult splits and tasks (e.g., protein structure prediction) to show a more pronounced difference between crammed and “fully trained” models, but we see these results as a promising sign that it is entirely possible to train a useful, expressive protein language model across the protein universe in 24 GPU hours and interrogate its performance on downstream tasks. Future work will address these limitations and extend the pLM cramming challenge to a more comprehensive evaluation of finetuning approaches and evaluation splits. We hope that our work will inspire efforts to cram pLMs, overhaul pLM training, and enable more fundamental investigations into language modeling for biological sequences.

## Acknowledgments and Disclosure of Funding

The authors acknowledge the entire Prescient Design team and the Antibody Engineering department at Genentech for providing helpful discussions and input that contributed to the research results reported within this paper. We acknowledge the Prescient Design Engineering team for providing HPC and consultation resources that contributed to the research results reported within this paper.

## Footnotes

1 https://github.com/prescient-design/lobster

## Notes

https://github.com/prescient-design/lobster

### Competing Interest Statement

All authors are or were employees of Genentech Inc., a member of the Roche Group, and may hold Roche stock or related interests.

